# Land-cover change in Cuba may favor biodiversity: An example using *Omphalea* (Angiosperma: Euphorbiaceae) and *Urania boisduvalii* (Lepidoptera: Uraniidae)

**DOI:** 10.1101/2023.02.17.529023

**Authors:** Claudia Nuñez-Penichet, Juan Maita-Chamba, Jorge Soberón

## Abstract

Changes in land cover directly affect biodiversity. Here we assessed land-cover change in Cuba in the past 35 years and analyzed how this change may affect the distribution of *Omphalea* plants and *Urania boisduvalii* moths. We analyzed the vegetation cover of the Cuban archipelago between 1985 and 2020. We used Google Earth Engine to classify two satellite image compositions into seven cover types: forest and shrubs, mangrove, soil without vegetation cover, wetlands, pine forest, agriculture, and water bodies. We considered three different areas for quantifications of land-cover change: (1) protected areas, (2) areas of potential distribution of *Omphalea*, and (3) areas of potential distribution of the plant within the protected areas. We found that the category of “forest and shrubs” have increased significantly in Cuba in the past 35 years and that most of the gained forest and shrub areas were agricultural land in the past. This same pattern was observed in the areas of potential distribution of *Omphalea*; whereas almost all cover types were mostly stables inside the protected areas. The transformation of agricultural areas into forest and shrubs could represent an interesting opportunity for biodiversity conservation in Cuba. Other detailed studies about biodiversity composition in areas of forest and shrubs gain would greatly benefit our understanding on the value of such areas for conservation.

## Introduction

Landscape degradation is one of the most important anthropogenic causes of biodiversity loss, driven mostly by changes in land use [1]. This is a problem for most species, but it may be especially important for those with a restricted distribution or those that depend on a specific climate or habitat for survival [2]. How habitat modifications are affecting species has been the focus of attention in several studies [3]. Those studies have been mainly focused on plants, birds, and mammals [4–6]. However, other groups like insects, are underrepresented [7], in spite of documented examples of extinctions due to habitat loss [8].

The neotropical genus *Urania* (Uraniidae) includes four species of diurnal moths, *U. sloanus* (endemic to Jamaica, presumed extinct; [9]), *U. fulgens* (from Mexico to northern Colombia), *U. leilus* (from southern Colombia to Bolivia), and *U. boisduvalii* (endemic to Cuba; [10]). These moths feed, during their larvae stages, exclusively on *Omphalea* (*Euphorbiaceae*) plants [9,11–13], which produce secondary metabolites as a defense mechanism against *Urania* herbivory [9]. These secondary compounds make the plants toxic for the moths, forcing them to move to a different host-plant population [14], therefore, *Urania* needs to have several patches of *Omphalea* available to guarantee its survival. Previous studies have also suggested that patches of *Omphalea* need to be structurally complex to support the moths’ larvae, as different larval stages use plants with different levels of maturity [15].

In Cuba, there are three species of *Omphalea* plants [9] mainly distributed in coastal and other karstic zones. *Omphalea trichotoma* (distributed in western and eastern Cuba) and *O. hypoleuca* (reported only from Viñales, Pinar del Río) are endemic [16], while *O. diandra* is widely distributed in the neotropics [9]. In this archipelago, these plants are present in areas that have been under pressure from the development of infrastructure for tourism and oil extraction activities [17] enhancing the importance of studying their populations and how they may be affected.

Given the direct dependance on host plant availability, *Urania* moths appear to be highly sensitive to changes in land use [9]. For instance, *U. sloanus* is now considered extinct due to habitat degradation [9]. However, no study has explored how distributional areas of these moths and plants are being affected over time due to land-cover change. Here, we aimed (1) to evaluate the change in land use in Cuba in the past 35 years, and (2) to analyze how these land use changes may affect the distribution of *Omphalea* plants and *Urania boisduvalii* moths.

## Methods

To evaluate land use change in Cuba (S1 Appendix 1), we analyzed vegetation cover of this archipelago between 1985 and 2020 using Landsat images and Google Earth Engine [18] for geospatial processing. Our study area was covered by sixteen scenes (paths: 10 to 17 and rows: 44 to 46) of the Landsat 5 ETM sensor (1985) and the Landsat 8 OLI/TIRS sensor (2020), with images at 30 meter resolution. To minimize temporal, spatial, and spectral varying scattering and absorbing effects of atmospheric gasses and aerosols, we used the collection 2 of Land Surface Reflectance [19]. Cloud-free images of Cuba were produced by using those with cloud cover ≤50% and a temporal filter including the years 1984-1988 for the period 1985 and the years 2020-2021 for 2020. Cloud and cloud shadows were masked using pixel quality attributes generated from the C Function of Mask (CFMASK) algorithm (QA_PIXEL Bitmask) for both sensors [20,21].

The resulting images (1985 and 2020) were classified into seven cover types to represent forest and shrubs (considered as suitable) and, mangrove, soil without vegetation cover, wetlands, pine forest, agriculture, and water bodies (considered as unsuitable areas for *Omphalea* plants; [22]). The classification of cover types was done by selecting regions of interest (ROI) for each of the categories on the Google Earth Engine. The number of ROIs selected for each category varied to represent how diverse they were (300 for forest and shrubs, 100 for mangrove, 50 for soil without vegetation cover, 100 for wetlands, 100 for pine forest, 100 for agriculture, and 100 for water bodies). All ROIs were selected manually based on visual interpretation of the respective Landsat images from Google Earth and were divided randomly into training (70%) and testing (30%).

We selected Random Forest as the classification algorithm as it has been described to be good at handling outliers and noisy datasets [23], have good performance with complex datasets [24] and higher accuracy than other algorithms [25,26], and its high processing speed [23]. The classified images were post-processed in QGIS [27] to correct misclassified pixels using a raster calculator and elevational considerations (S1 Appendix 2). We assessed the accuracy of classifications via: 1) overall accuracy (OA) which is calculated dividing the total number of correctly classified values by the total number of values [28,29], and 2) kappa coefficient [30,31] for measuring the agreement between classification and truth values. A kappa value of 1 represents perfect agreement, while a value of 0 represents no agreement. These steps were done on the Google Earth Engine platform, using the resulting images from the post-processing analyses and 100 new validation ROIs created randomly for each of the considered cover types but water bodies in which we used only 50.

The resulting classified images for Cuba were masked to: (1) terrestrial protected areas [32]; (2) potential distribution areas of *Omphalea* (from [22], hereafter OD; and (3) potential distribution areas of the plant within protected areas. We quantified the percentage of each of the seven cover types and their changes between periods for all Cuba and in each of the masked areas (areas of interest). All these analyses were done in R [33] using the packages rgdal [34] and raster [35]. All the code needed to reproduce these analyses is openly available at https://github.com/claununez/Land-coverChangeCuba. All the data needed is openly available at https://figshare.com/articles/dataset/Data_for_the_manuscript_Land-cover_change_in_Cuba_may_favor_biodiversity_An_example_using_Omphalea_plants_and_Urania_boisduvalii/21779129.

## Results

We obtained an overall accuracy (OA) of 92.7% and 87.6% for 1985 and 2020 classifications, with a Kappa coe□cient of 0.91 and 0.85, respectively. The values of OA for each land cover type ranged from 80% (pine forest) to 100% (agriculture) in 1985, and from 71% (wetlands) to 99% (forest and shrubs) in 2020 (S1 Table).

We found that, in the past 35 years, forest and shrub areas have increased across the entire Cuban territory (from 23.5% in 1985 to 38.89% in 2020; Table 1). Agricultural lands and soils without vegetation cover, on the other hand, showed a reduction from 50.18% to 39.12% and from 6.03% to 1.85%, respectively (Table 1; Fig 1). The other four types of cover considered (mangrove, wetland, pine forest, and water bodies) were present in similar proportions in the two scenarios studied (Table 1). Most areas that changed from one cover type to another were located inland, especially in areas of low elevation (Fig 1; S1 Appendix 1).

**Table 1.** Percentage of area classified as one of the seven classification types considered for the years 1985 and 2020. *Omphalea* includes all the species of *Omphalea* genus distributed in Cuba combined and the areas of potential distribution of *Omphalea* is referring to these species’ potential distribution in Cuba. The protected areas in Cuba are referring to the terrestrial protected areas only.

| Classification type | Cuba |  | Protected areas in Cuba |  | Areas of potential distribution of *Omphalea* |  | Areas of potential distribution of *Omphalea* inside protected areas |  |
| --- | --- | --- | --- | --- | --- | --- | --- | --- |
|  | 1985 | 2020 | 1985 | 2020 | 1985 | 2020 | 1985 | 2020 |
| Forest and shrubs | 23.50 | 38.89 | 33.80 | 36.42 | 57.67 | 73.26 | 71.24 | 74.96 |
| Mangrove | 4.03 | 4.16 | 13.05 | 12.82 | 1.66 | 1.91 | 3.47 | 3.86 |
| Soil without vegetation cover | 6.03 | 1.85 | 4.62 | 0.88 | 2.61 | 0.55 | 2.01 | 0.33 |
| Wetland | 5.84 | 6.08 | 17.13 | 19.30 | 2.68 | 2.02 | 4.34 | 3.86 |
| Pine forest | 2.28 | 1.24 | 2.43 | 2.07 | 5.60 | 3.44 | 6.42 | 5.46 |
| Agriculture | 50.18 | 39.12 | 7.92 | 6.86 | 26.92 | 15.64 | 8.04 | 6.70 |
| Water bodies | 8.14 | 8.65 | 21.05 | 21.65 | 2.86 | 3.19 | 4.48 | 4.82 |

**Figure 1.**
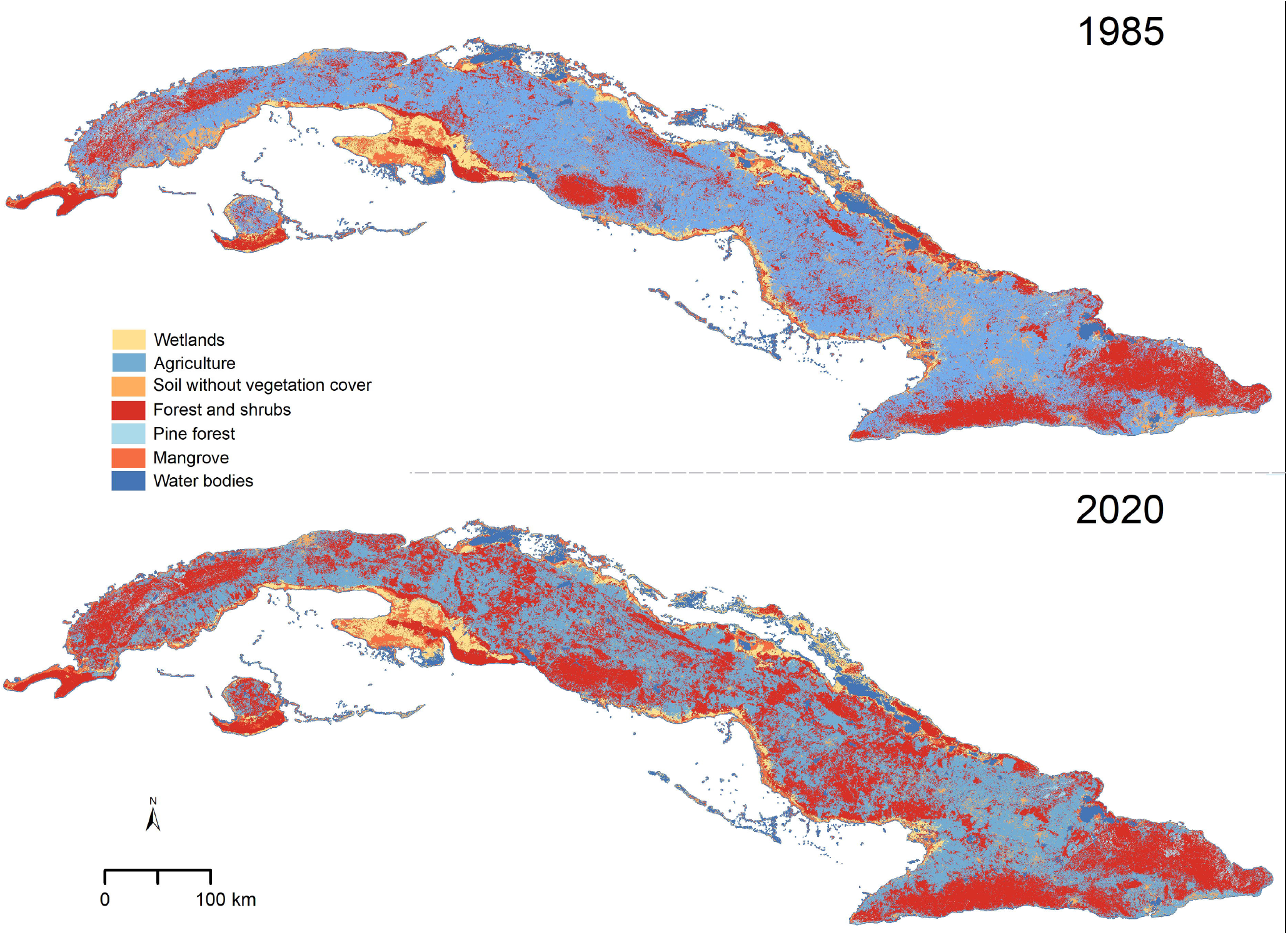
Vegetation types in Cuba for the years 1985 and 2020.

In the areas of potential distribution of *Omphalea* plants, we found a similar pattern to the one described above for Cuba (Table 1). Most of these changes were concentrated in central Cuba, whereas stable areas were mostly in Guanahacabibes (westernmost part of Cuba), the southern part of Isla de la Juventud, and highlands (S1 Appendix 1; S1 Figure).

The percentage of area of each cover type was similar between 1985 and 2020 inside the protected areas in Cuba and in the OD that were inside the protected areas (Table 1, S2 Figure, S3 Figure). The only exception was soils without vegetation cover, which decreased from 4.62% to 0.88% inside protected areas in Cuba, and from 2.01% to 0.33% in the OD inside protected areas (Table 1, S2 Figure, S3 Figure).

We found that only 19.16% of Cuba classified as forest and shrubs in 2020, was classified under the same category in 1985 (Table 2). The low stability of this cover type was also found when quantifying in the other areas of interest, with 29.68% of the protected areas in Cuba, 52.52% of the OD, and 64.49% of the OD inside protected areas found to be forest and shrubs in both time periods analyzed (Table 2). Most of the forest and shrub areas gained in Cuba in 2020 were agricultural lands in the past (Table 2). The proportion of change from agricultural land to forest and shrubs was also high in the OD and the other areas of interest (Table 2). The agricultural lands that changed to forest and shrubs by 2020 were distributed in small patches across the Cuban archipelago and were less predominant in the eastern region (Fig 2). The areas that changed to forest and shrubs in OD were also small patches, but their spatial distribution helped increase the connectivity among larger stable forest and shrubs areas (Fig 2, Table 2).

**Table 2.** Percentage of change between the classification types considered, during the period 1985 - 2020. *Omphalea* includes all the species of the genus *Omphalea* distributed in Cuba combined and the areas of potential distribution of *Omphalea* is referring to these species’ potential distribution in Cuba.

| Type of change | Cuba | Protected areas in Cuba | Areas of potential distribution of *Omphalea* | Areas of potential distribution of *Omphalea* inside protected areas |
| --- | --- | --- | --- | --- |
| Stable forest and shrubs | 19.16 | 29.68 | 52.52 | 64.49 |
| Mangrove to forest and shrubs | 0.19 | 0.65 | 0.30 | 0.80 |
| Soil without vegetation cover to forest and shrubs | 0.92 | 0.12 | 0.74 | 0.10 |
| Wetlands to forest and shrubs | 0.75 | 1.31 | 1.02 | 1.34 |
| Pine forest to forest and shrubs | 1.41 | 1.45 | 3.66 | 4.03 |
| Agriculture to forest and shrubs | 16.45 | 3.21 | 15.46 | 4.20 |
| Water bodies to forest and shrubs | 0.01 | 0.01 | 0.01 | 0.02 |
| Stable mangrove | 2.88 | 8.94 | 0.40 | 1.86 |
| Forest and shrubs to mangrove | 0.35 | 1.08 | 0.40 | 1.06 |
| Soil without vegetation cover to mangrove | 0.03 | 0.07 | 0.02 | 0.02 |
| Wetlands to mangrove | 0.70 | 2.28 | 0.38 | 0.77 |
| Pine forest to mangrove | 0.01 | 0.03 | 0.02 | 0.05 |
| Agriculture to mangrove | 0.06 | 0.09 | 0.05 | 0.05 |
| Water bodies to mangrove | 0.13 | 0.34 | 0.04 | 0.05 |
| Stable soil without vegetation cover | 1.05 | 0.58 | 0.35 | 0.23 |
| Forest and shrubs to soil without vegetation cover | 0.03 | 0.04 | 0.03 | 0.04 |
| Mangrove to soil without vegetation cover | 0.00 | 0.00 | 0.00 | 0.00 |
| Wetlands to soil without vegetation cover | 0.06 | 0.17 | 0.02 | 0.04 |
| Pine forest to soil without vegetation cover | 0.00 | 0.00 | 0.00 | 0.00 |
| Agriculture to soil without vegetation cover | 0.70 | 0.07 | 0.15 | 0.03 |
| Water bodies to soil without vegetation cover | 0.00 | 0.01 | 0.00 | 0.00 |
| Stable wetlands | 3.77 | 12.23 | 1.00 | 1.76 |
| Forest and shrubs to wetlands | 0.23 | 0.49 | 0.13 | 0.20 |
| Mangrove to wetlands | 0.80 | 2.98 | 0.28 | 0.65 |
| Soil without vegetation cover to wetlands | 0.81 | 2.67 | 0.44 | 0.99 |
| Pine forest to wetlands | 0.00 | 0.00 | 0.00 | 0.00 |
| Agriculture to wetlands | 0.29 | 0.27 | 0.09 | 0.07 |
| Water bodies to wetlands | 0.19 | 0.66 | 0.08 | 0.18 |
| Stable pine forest | 0.46 | 0.69 | 1.27 | 1.74 |
| Forest and shrubs to pine forest | 0.47 | 1.12 | 1.63 | 3.22 |
| Mangrove to pine forest | 0.00 | 0.00 | 0.00 | 0.00 |
| Soil without vegetation cover to pine forest | 0.01 | 0.00 | 0.00 | 0.00 |
| Wetlands to pine forest | 0.00 | 0.00 | 0.00 | 0.00 |
| Agriculture to pine forest | 0.30 | 0.26 | 0.55 | 0.50 |
| Water bodies to pine forest | 0.00 | 0.00 | 0.01 | 0.01 |
| Stable agriculture | 32.12 | 4.02 | 10.68 | 3.19 |
| Forest and shrubs to agriculture | 3.15 | 1.31 | 3.24 | 2.23 |
| Mangrove to agriculture | 0.05 | 0.13 | 0.04 | 0.07 |
| Soil without vegetation cover to agriculture | 3.01 | 0.47 | 0.93 | 0.32 |
| Wetlands to agriculture | 0.40 | 0.66 | 0.19 | 0.28 |
| Pine forest to agriculture | 0.35 | 0.26 | 0.65 | 0.59 |
| Water bodies to agriculture | 0.04 | 0.02 | 0.02 | 0.01 |
| Stable water bodies | 7.78 | 20.01 | 2.72 | 4.21 |
| Forest and shrubs to water bodies | 0.12 | 0.08 | 0.08 | 0.02 |
| Mangrove to water bodies | 0.11 | 0.36 | 0.04 | 0.08 |
| Soil without vegetation cover to water bodies | 0.20 | 0.71 | 0.14 | 0.35 |
| Wetlands to water bodies | 0.16 | 0.47 | 0.08 | 0.15 |
| Pine forest to water bodies | 0.05 | 0.00 | 0.03 | 0.00 |
| Agriculture to water bodies | 0.24 | 0.02 | 0.11 | 0.00 |

**Figure 2.**
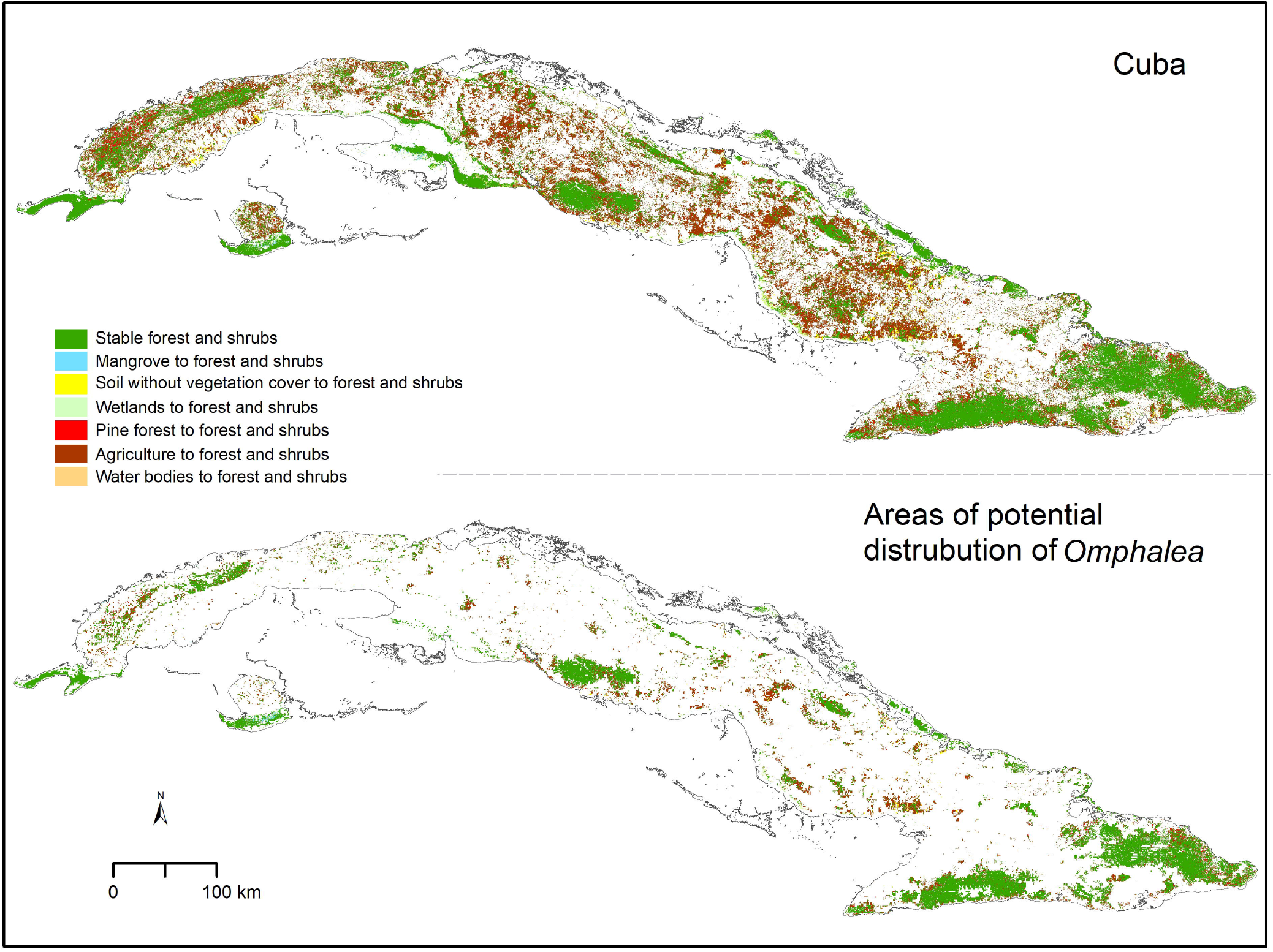
Forest and shrubs change. *Omphalea* includes all the species of *Omphalea* genus distributed in Cuba combined and the areas of potential distribution of *Omphalea* is referring to these species’ potential distribution in Cuba.

## Discussion

The general global trend in forest cover is towards loss [36]. However, we found that in Cuba, forest and shrubs (i.e., suitable vegetation cover for *Omphalea*) have increased considerably in the past 35 years, mostly replacing agricultural lands (Fig 1, Table 1). This result may be due to the fact that in Cuba, before 1990, extensive areas were dedicated to growing sugarcane, especially in the center of the island (S1 Appendix 1), but after 1990, the intensity of sugarcane production decreased [37,38]. Our results concur with [39] who reported that over half of agricultural land in Cuba is currently unused. Although the trend we report is opposite to the general global pattern, it is an example of how land use change is a complex phenomenon resulting from multiple causes, and difficult to predict [40].

Forest and shrub cover increases within areas of potential distribution of *Omphalea* (Table 1; S1 Figure) may in fact represent an advantage for populations of these plants as these areas are known to be climatically suitable [22]. Given the obligated relationship between *Urania boisduvalii* and *Omphalea* plants, which also drives the moth’s dispersal patterns in the country (Nuñez-Penichet et al., in prep), these gained areas could indirectly benefit the moth. This is because the stability and increment of this type of land cover could be increasing connectivity among fragmented patches with plant populations (Fig 2; S1 Figure). In fact, several of the stable and gained shrub and forest patches are located in the potential migratory paths of *U. boisduvalii* identified in a previous study [41].

On the other hand, the increment of forest and shrubs in the Cuban territory does not necessarily imply an increase of *Omphalea* plants or an increment in favorable conditions for *Urania* moths. Species distributions depend on the presence of suitable abiotic conditions, accessibility, and biotic interactions [42]. In Cuba, a significant portion of unused agricultural lands have been reported to be covered with invasive species (e.g., *Dichrostachys cinerea*, [39,43]; and *Vachellia farnesiana*, [44], which causes a displacement of native plant communities [45]. In the last decades, several studies have focused on testing the potential of *D. cinerea* high-quality biomass to produce a sustainable biofuel [39,46] as an alternative source of energy, and therefore little an effort is targeted to control this invasive plant. For this reason, performing fieldwork in the gained forest and shrubs areas will be necessary to assess whether they can actually benefit the conservation of *Omphalea* plants and *Urania* moths.

When we focused on potential distribution areas for *Omphalea* inside protected areas, we detected that stability was high for all cover types explored, except for soils without vegetation cover (S2 Figure, S3 Figure). This concurs with what has been previously reported for 10 National Parks in Cuba [47]. However, interpretations about the conservation status of natural vegetation inside protected areas should be done cautiously, as given the goal of our study, we included all types of forests and shrubs in a single category. Therefore, we are not accounting for changes at a finer detail, and forest and shrub gains do not necessarily represent positive rates of change for primary vegetation.

Here we are presenting results that suggest the distribution of *Omphalea* plants and for *Urania* moths in Cuba may be increasing, since we find that the areas with suitable conditions for these species have been increasing considerably. Moreover, our results suggest that land-use changes in Cuba may favor not only *Urania* and *Omphalea* but other species that live in forests and shrubs. However, the presence of invasive species in those gained forest and shrubs areas should be monitored in the field as they may represent a serious threat to native biodiversity.

Changes in land use in Cuba during the last 35 years may represent a significant opportunity for biodiversity conservation and sustainable management of its natural vegetation. However, further studies that allow understanding the community composition and structure in such forests and shrubs are needed to assess the value of these lands for biodiversity conservation in the country.

## Acknowledgments

We thank Marlon E. Cobos for his useful comments on the manuscript.

## Supporting information captions

S1 Table. Values of the overall accuracy of the classification, for the years 1985 and 2020, for each land cover type considered in this study.

S1 Figure. Vegetation types in the areas of potential distribution of *Omphalea* plants in Cuba for the years 1985 and 2020. *Omphalea* includes all the species of *Omphalea* genus distributed in Cuba combined.

S2 Figure. Vegetation types in the terrestrial protected areas of Cuba for the years 1985 and 2020.

S3 Figure. Cover types in the areas of potential distribution of *Omphalea* plants in Cuba that are inside the terrestrial protected areas. *Omphalea* includes all the species of *Omphalea* genus distributed in Cuba combined and the areas of potential distribution of *Omphalea* is referring to these species’ potential distribution in Cuba.

S1 Appendix 1. Description of the area of study.

S1 Appendix 2. Elevational considerations used for post-processing the classified images to correct misclassified pixels.

