## Supplementary information for "Land-cover change in Cuba may favor biodiversity: An example using *Omphalea* (Angiosperma: Euphorbiaceae) and *Urania boisduvalii* (Lepidoptera: Uraniidae)"

**Supplementary materials for the manuscript “Land-cover change in Cuba may favor biodiversity: An example using *Omphalea* (Angiosperma: Euphorbiaceae) plants and *Urania boisduvalii* (Lepidoptera: Uraniidae)”**

Claudia Nuñez-Penichet^1*^, Juan Maita-Chamba^2^, and Jorge Soberón^1^

^1^Biodiversity Institute and Department of Ecology & Evolutionary Biology, University of Kansas.

^2^Carrera de Ingeniería Forestal, Centro de Investigaciones Tropicales del Ambiente y Biodiversidad, Universidad Nacional de Loja, Loja 110111, Ecuador

^*^Corresponding Author: Claudia Nuñez-Penichet

Biodiversity Institute and Department of Ecology & Evolutionary Biology, University of Kansas, 1345 Jayhawk Blvd., Lawrence, Kansas 66045, USA.

S1 Table. Values of the overall accuracy of the classification, for the years 1985 and 2020, for each land cover type considered in this study.

| Cover type | Overall accuracy (%) | |
| --- | --- | --- |
|  | 1985 | 2020 |
| Forest and shrubs | 98 | 99 |
| Mangrove | 91 | 95 |
| Soil without vegetation cover | 83 | 89 |
| Wetlands | 92 | 71 |
| Pine forest | 80 | 72 |
| Agriculture | 100 | 95 |
| Water bodies | 97 | 98 |

S1 Figure. Vegetation types in the areas of potential distribution of *Omphalea* plants in Cuba for the years 1985 and 2020. *Omphalea* includes all the species of *Omphalea* genus distributed in Cuba combined.

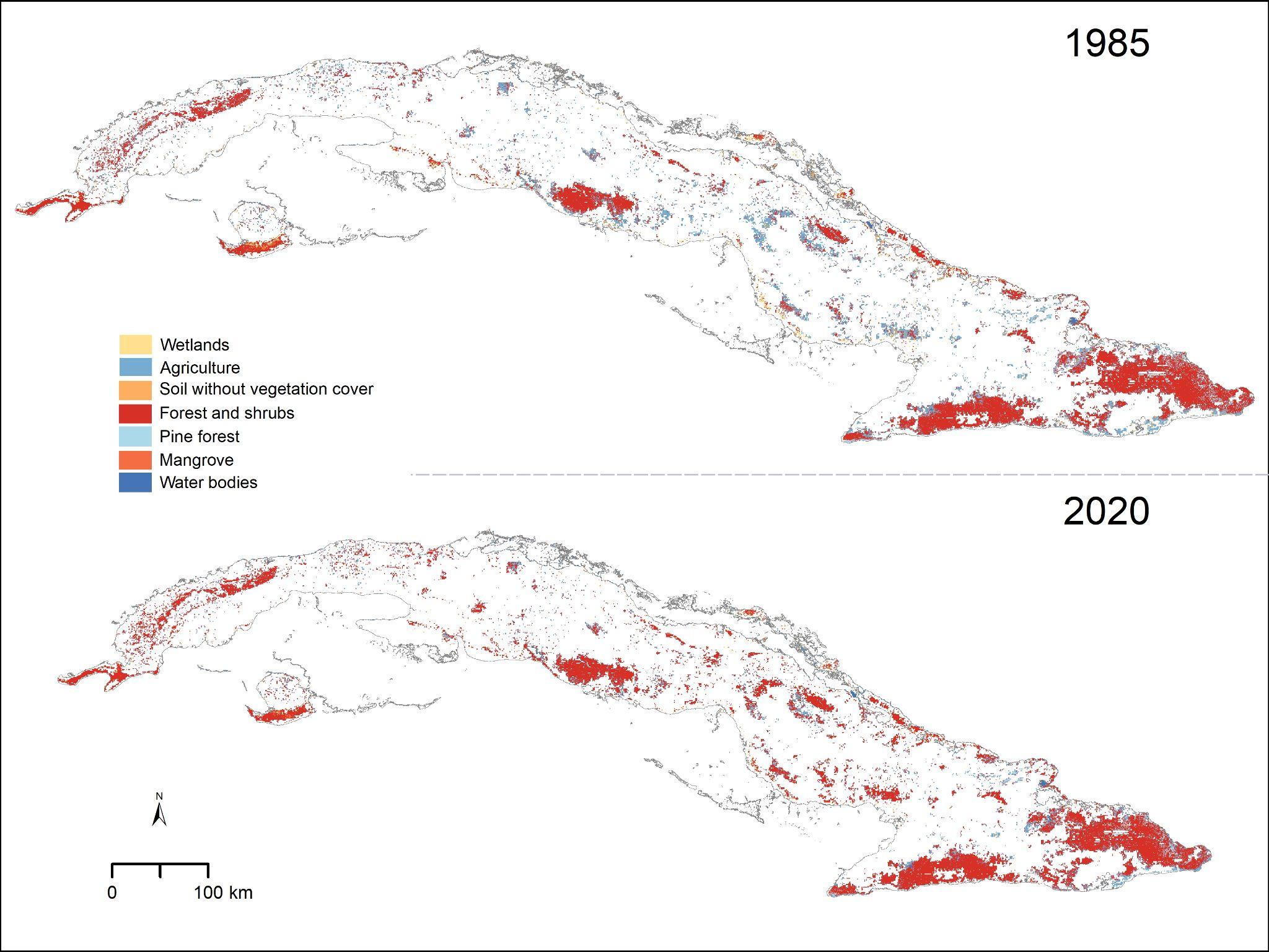

S2 Figure. Vegetation types in the terrestrial protected areas of Cuba for the years 1985 and 2020.
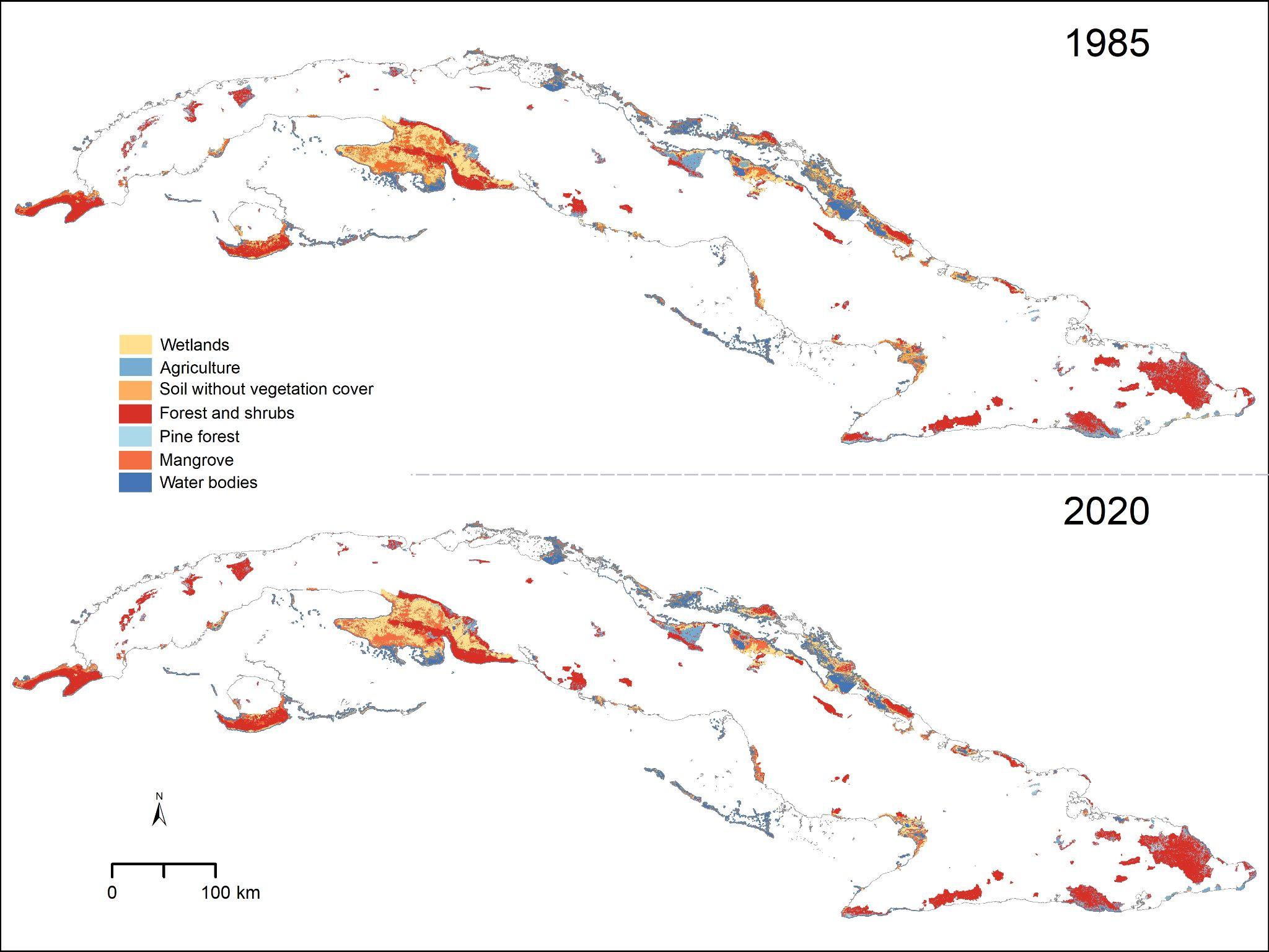

S3 Figure. Cover types in the areas of potential distribution of *Omphalea* plants in Cuba that are inside the terrestrial protected areas. *Omphalea* includes all the species of *Omphalea* genus distributed in Cuba combined and the areas of potential distribution of *Omphalea* is referring to these species’ potential distribution in Cuba.
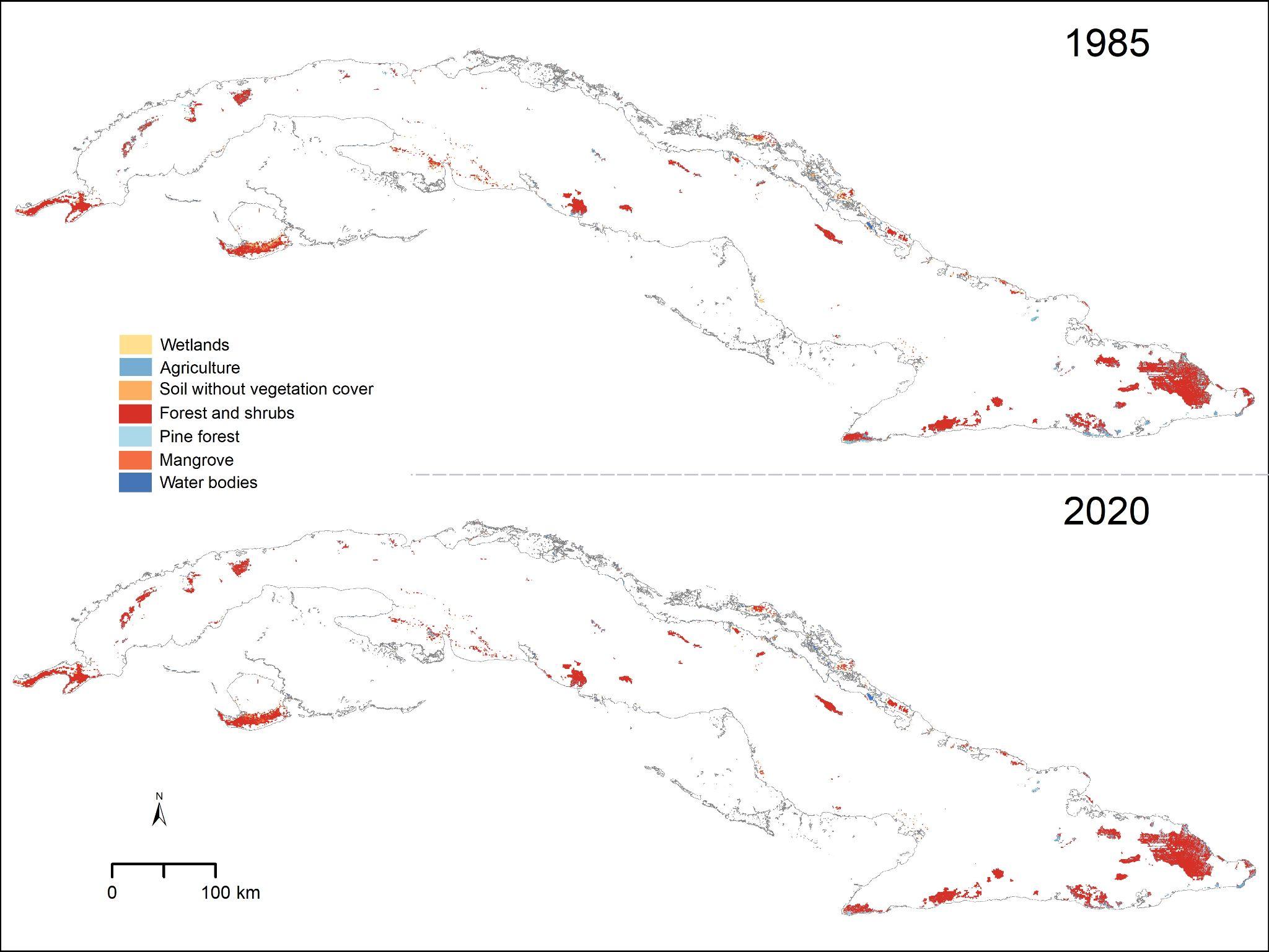

S1 Appendix 1. Description of the area of study.

Cuban archipelago is located on the northeast side of the Caribbean Sea (-84.993 – -74.111 longitude, 19.837 – 23.233 latitude), is part of the Greater Antilles, and has a surface of approximately 110922 km2 (Gutiérrez and Rivero, 1997; Fig. 1). Cuba’s landscape is formed mostly by lowlands with few elevations (Díaz, 1989; Fig. 1). The neotropical location of this archipelago, which determine its temperature and precipitation, together with the tectonic movements and volcanic activity from the past (Formel, 1989), influence the soil types of Cuba as well as its vegetation (Gutiérrez and Rivero, 1997).

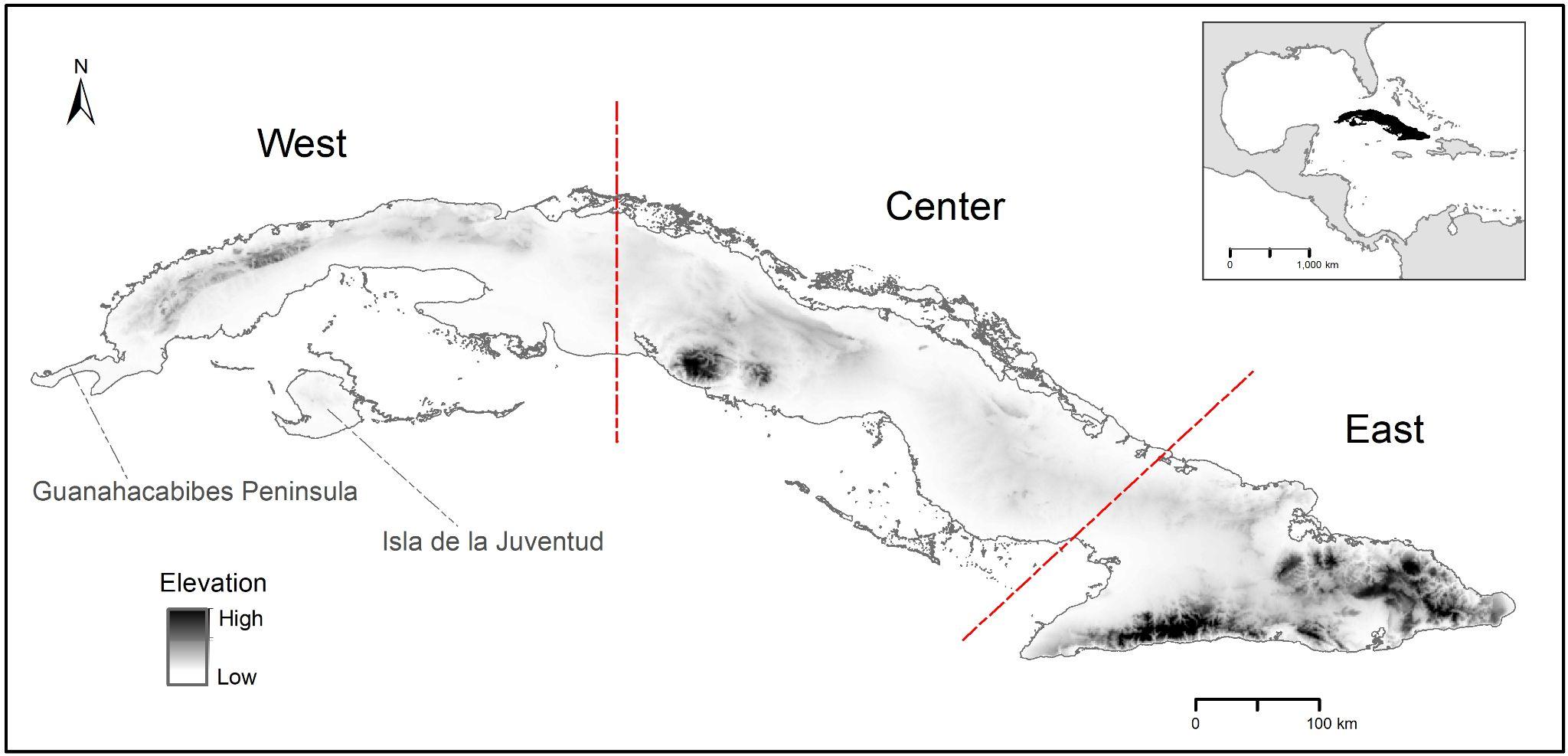

Figure 1. Study area.

S1 Appendix 2. Elevational considerations used for post-processing the classified images to correct misclassified pixels.

**Reclassification of pixels on 1985 land cover map using elevational considerations**

1. Mangrove pixels ⋝ 20 m above sea level change to pinares
2. Wetland pixels ⋝ 11 m above sea level change to agriculture
3. Pinares pixels ⋜ 20 m above sea level change to wetlands
4. Agriculture land pixels ⋜ 4 m above sea level change to wetlands
5. Water bodies pixels ⋝ 200 m above sea level change to forest

**Reclassification of pixels on 2020 land cover map using elevational considerations**

1. Mangrove pixels ⋝ 20 m above sea level change to pinares
2. Wetland pixels ⋝ 11 m above sea level change to agriculture
3. Pinares pixels ⋜ 22 m above sea level change to mangroves
4. Agriculture land pixels ⋜ 4 m above sea level change to wetlands
5. Water bodies pixels ⋝ 300 m above sea level change to forest
